# A linear DNA vaccine candidate encoding the SARS-CoV-2 Receptor Binding Domain elicits protective immunity in domestic cats

**DOI:** 10.1101/2022.07.20.500860

**Authors:** Antonella Conforti, Elisa Sanchez, Erika Salvatori, Lucia Lione, Mirco Compagnone, Eleonora Pinto, Fabio Palombo, Yuhua Sun, Brian Viscount, James Hayward, Clay Shorrock, Diego G. Diel, Joseph A. Impellizeri, Luigi Aurisicchio

## Abstract

Since its first detection in China in late 2019, SARS-CoV-2, the etiologic agent of COVID-19 pandemic, has infected a wide range of animal species, especially mammals, all over the world. Indeed, as reported by the American Veterinary Medical Association, besides human-to-human transmission, human-to-animal transmission has been observed in some wild animals and pets, especially in cats. With animal models as an invaluable tool in the study of infectious diseases combined with the fact that the intermediate animal source of SARS-CoV-2 is still unknown, researchers have demonstrated that cats are permissive to COVID-19 and are susceptible to airborne infections. Given the high transmissibility potential of SARS-CoV-2 to different host species and the close contact between humans and animals, it is crucial to find mechanisms to prevent the transmission chain and reduce the risk of spillover to susceptible species. Here, we show results from a randomized Phase I/II clinical study conducted in domestic cats to assess safety and immunogenicity of a linear DNA (“linDNA”) vaccine encoding the RBD domain of SARS-CoV-2. No significant adverse events occurred and both RBD-specific binding/neutralizing antibodies and T cells were detected. These findings demonstrate the safety and immunogenicity of a genetic vaccine against COVID-19 administered to cats and strongly support the development of vaccines for preventing viral spread in susceptible species, especially those in close contact with humans.

## INTRODUCTION

Since the first detection and initial outbreak of coronavirus disease (COVID-19) pandemic in late 2019, caused by the severe acute respiratory syndrome coronavirus 2 (SARS-CoV-2), epidemiological investigations suggested the animal origin of the infectious agent, possibly from bat^1^. Via a yet unknown intermediate animal host, the virus quickly adapted to humans and human-to-human transmission rapidly became the principal and immediate source of virus spread. The wide and accelerated distribution in the human population accompanied by specific risk factors, such as close interaction between infected humans and susceptible animal species and the suitability of animal host population for viral infection, has considerably increased the risk of human-to-animal transmission, i.e. reverse zoonoses^2^.

Due to its zoonotic origin, SARS-CoV-2 may also be relevant to animals. Thus, to evaluate the host range of the virus and to assess the risk to act as potential animal reservoir, a large number of different animal species were experimentally infected with SARS-CoV-2 or monitored in the field in the last months^3,4^. To date, SARS-CoV-2 infections associated with zoonotic transmission have been identified in domestic cats and dogs, farmed wildlife such as minks and ferrets, zoo animals such as tigers and lions, and free-ranging wildlife such as white-tailed deer^5,6,7,8^. The reverse zoonosis phenomenon may become a potential threat not only for animal health itself but also for human health, given that infected animals can become a virus reservoir, from which reintroduction into humans can occur^9^.

Vaccines are one of the most effective tools for preventing and controlling infectious diseases, such as COVID-19. Since the outbreak of the COVID-19 pandemic, a number of safe and effective vaccines have been developed for use in humans, (including recombinant protein vaccines, adenoviral vector-based and nucleic acid vaccines) and animals (recombinant protein vaccine). Among the nucleic acid-based vaccines, DNA-based platforms show great potential in terms of safety and ease of production^10^. Recently, we showed that COVID-eVax, a DNA-based vaccine encoding the receptor-binding domain (RBD) of SARS-CoV-2 spike protein (S) and delivered by intramuscular electro-gene-transfer (EGT), is a highly efficient vaccination platform capable of inducing robust, protective neutralizing antibody and T cell responses in a variety of animal models^11^. Moreover, COVID-eVax has demonstrated its safety and immunogenicity in a first-in-human clinical trial (manuscript submitted). Besides this, new cell-free manufacturing DNA platforms, such as doggybone DNA and PCR-based amplicons such as linDNA, have proven to be stable and immunogenic in preclinical models, thus allowing a highly scalable manufacturing workflow in perspective of massive vaccination for future pandemics^12,13^. Therefore, in order to ensure preparedness in response to increasing reports of COVID-19 in companion animals, we evaluated the safety and immunogenicity of a PCR-based linDNA vaccine, encoding RBD, delivered by EGT (DNA-EGT) in domestic cats. We show that immunization of cats with the vaccine candidate results in induction of RBD-specific T cell response and neutralizing antibodies against SARS-CoV-2 and its variants.

## RESULTS

### Manufacturing of linDNA vaccine

PCR primers were designed to amplify the expression cassette encoding the RBD domain of SARS-CoV-2 Spike protein contained in the pTK1A-TPA-RBD plasmid. The first five bases from the 5’ terminal end of both forward and reverse primers were modified with Phosphorothioate Bonds - a sulfur atom substitutes the non-bridging oxygen in the phosphate backbone of an oligonucleotide - to increase linDNA stability^14^. PCR amplifications were performed using a large-scale PCR system with Q5™ polymerase (5.3×10^−7^ approximate error/bp; New England BioLabs, USA). The PCR mixture consisted of 1X buffer, 0.8mM dNTP, 0.5μM of forward and reverse primers, 1U/100ul of Q5 polymerase, and 40ng/mL of plasmid template. The PCR reactions were subjected to initial denaturation of 98°C for 45 seconds, and subsequent 30 cycles of 97°C for 80 seconds and 70°C for 4.5 minutes, followed by final extension at 72°C for 3 minutes. Upon completion of the PCR reaction the linDNA expression cassettes were purified and ready for use in the transfection of target cells for expression of the RBD domain of SARS-CoV-2 Spike protein. The purity and integrity of the linDNA expression cassettes as measured by HPLC are shown in **Fig. 1A**.

**Figure 1.**
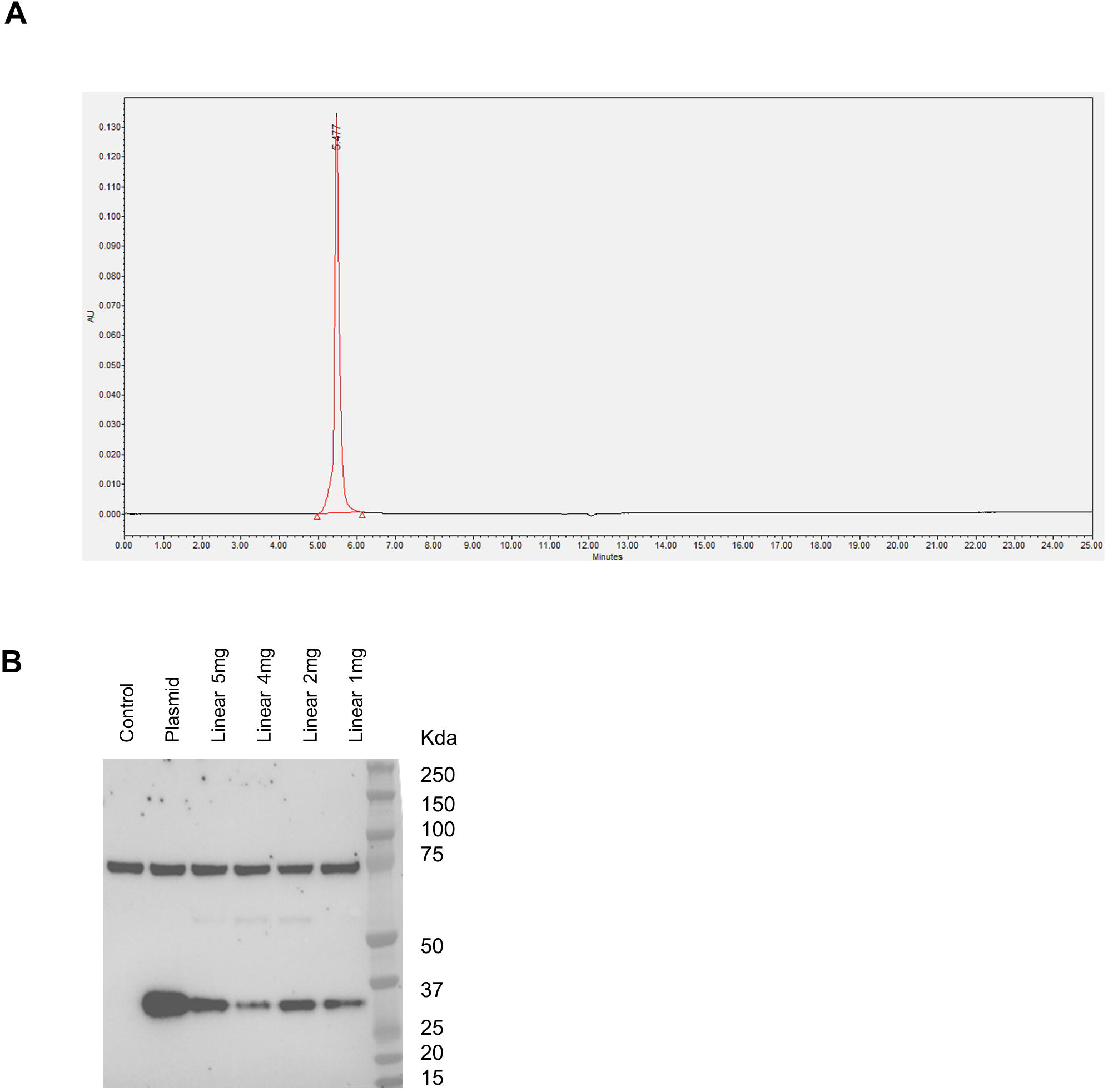
Characterization and expression of linDNA vaccine. **(A)** HPLC chromatogram of linDNA. **(B)** Western blot analysis of both plasmid and linDNA vaccine encoding RBD from SARS-CoV-2 Spike protein, after transfection in HEK293 cells. Forty-eight hours after transfection, cell lysates were resolved on a gel and blotted with a polyclonal SARS-CoV-2 Spike S1 Subunit antibody. Cells transfected with empty plasmid vector were used as negative control (control).

### In vitro expression of linDNA vaccine

A plasmid DNA encoding the RBD domain of SARS-CoV-2 Spike protein (pTK1A-TPA-RBD) was used as template for the PCR-based linDNA amplicon expression vector synthesis. In order to promote protein secretion, we introduced a tissue plasminogen activator (tPA) leader sequence. As shown in **Fig.1B**, Western blot analyses confirmed the expression of RBD in cell lysates from HEK293 cells transiently transfected with either 5μg of plasmid DNA or decreasing doses of PCR-based linDNA amplicon (from 5 to 1 μg).

### linDNA COVID-19 vaccine immunogenicity in mice

In order to assess the in vivo immunogenicity of the linDNA COVID-19 vaccine, electroporation of either plasmid or linDNA encoding RBD domain of SARS-CoV-2 Spike protein in the skeletal muscle of BALB/c mice was performed. The vaccination protocol consisted of the injection of 20 μg of plasmid DNA (10 μg per quadriceps) or equimolar dose of linDNA (4,3 μg per quadriceps) into 6-week-old BALB/c mice at day 0 (prime) and day 28 (boost) with sacrifice at day 38 (**Fig. 2A**). The humoral response in the sera of vaccinated mice was evaluated by measuring anti-RBD IgG titers by ELISA at day 14 (prime) and at day 38 (boost) (**Fig. 2B**). At day 14 all mice showed detectable anti-RBD IgG antibodies, and their levels increased at day 38. Of note, anti-RBD IgG titers induced by plasmid DNA were significantly higher than those elicited by linDNA. Next, we sought to compare the T cell response elicited by either plasmid or linDNA RBD vaccine by ELISpot assay. To this end, we used a peptide pool covering the RBD domain of the Spike protein to stimulate splenocytes collected from BALB/c mice at day 38 after vaccination. Different from humoral response, RBD-specific T cell response was significantly higher in linDNA vaccinated group (**Fig.2C**). These results clearly demonstrated that linDNA vaccine is able to elicit both humoral and cellular immune response specific for SARS-CoV-2 RBD in a preclinical model.

**Figure 2.**
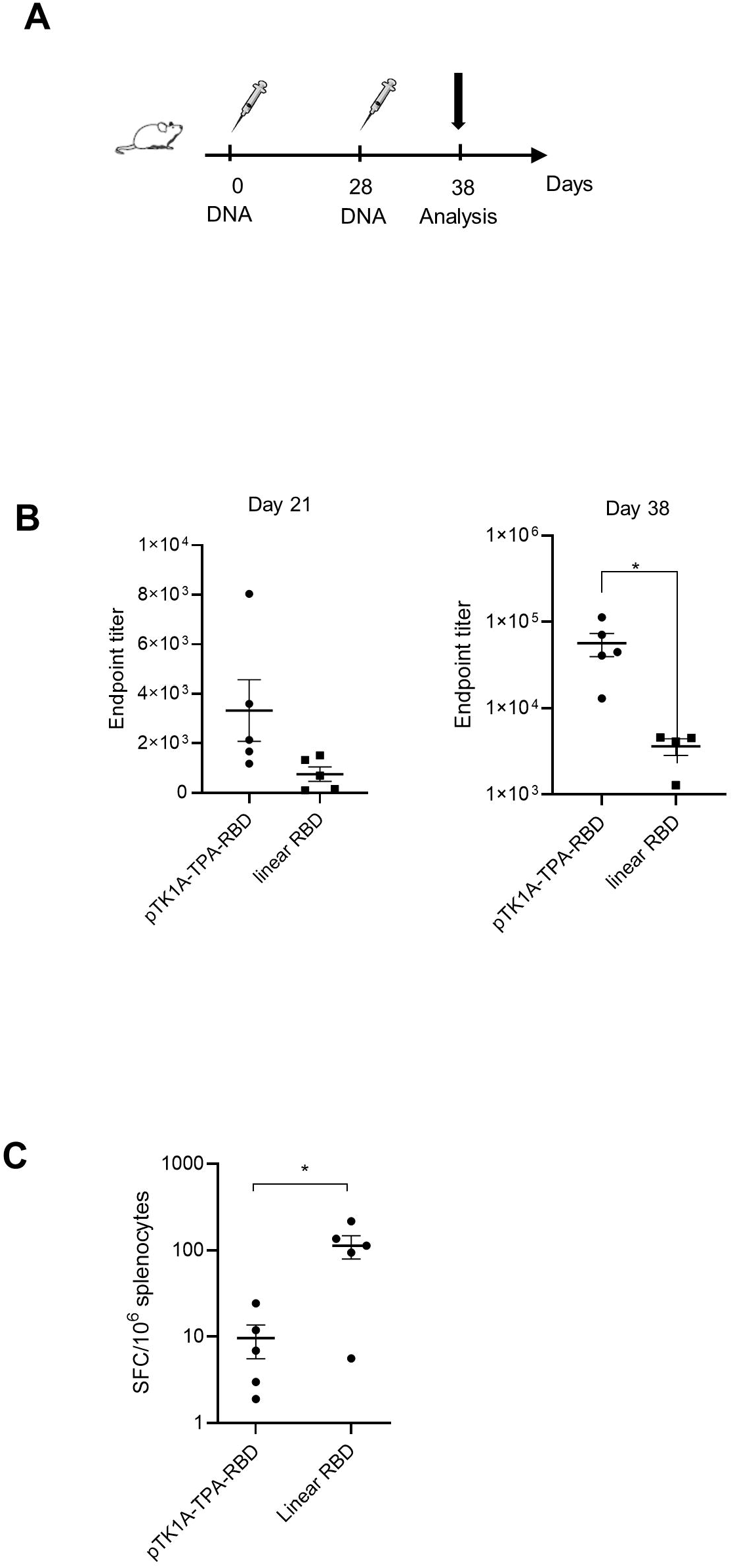
In vivo immunogenicity of linDNA vaccine. **(A)** Schematic representation of experimental setup. 6-week-old Balb/c mice (5 mice/group) were injected, by means of i.m. electroporation, with 20 μg of plasmid DNA (10 μg per quadriceps) or equimolar dose of linDNA (4,3 μg per quadriceps) at day 0 (prime) and day 28 (boost) with sacrifice at day 38. **(B)** Anti-RBD IgG endpoint titer measured by ELISA assay performed on sera collected at day 21 (after prime) and 38 (after boost) from mice vaccinated with either plasmid DNA or linDNA. **(C)** IFNγ-producing T cells measured by ELISpot assay performed on splenocytes collected from mice vaccinated with either plasmid DNA or linDNA and culled at day 38. Significance was determined using Mann-Whitney test, *p<0,05 **p< 0.01.

### Feline study design and safety

To assess linDNA vaccine efficacy in an animal species naturally susceptible to SARS-CoV-2 infection, a total of 11 domestic cats were enrolled in a randomized Phase I/II clinical study (**Table I**). The vaccination regimen is shown in **Fig. 3A** and consisted of two injections (days 0 and 28) of 1mg linDNA intramuscularly, followed immediately by co-localised intramuscular electroporation (by means of Vet-ePorator™) into the tibialis cranialis region of each rear leg, as previously described^15^. The profile of electroporation parameters was checked at every DNA administration in real time and data were stored into the Vet-ePorator™ archive for each single animal. The procedure lasted less than 20 seconds and each patient recovered uneventfully each time.

**Table 1.** Demographics

| ID | Breed | Weight (Kg) | Gender | Age (months) |
| --- | --- | --- | --- | --- |
| 1 | Domestic Shorthair | 5.9 | FS | 87 |
| 2 | Ragdoll | 4.25 | MN | 166 |
| 3 | Domestic Shorthair | 8.9 | MN | 61 |
| 4 | Domestic Shorthair | 3.4 | MN | 120 |
| 5 | Norwegian Forest Cat | 6.5 | MN | 73 |
| 6 | Domestic Shorthair | 4.55 | MN | 37 |
| 7 | Domestic Shorthair | 6.4 | MN | 167 |
| 8 | Domestic<br>Shorthair | 4.65 | MN | 31 |
| 9 | Domestic<br>Shorthair | 8.7 | MN | 46 |
| 10 | Domestic<br>Shorthair | 5.7 | FS | 5.7 |
| 11 | Domestic<br>Shorthair | 5.3 | MN | 5.3 |

MN, Male Neutered; FS, Female Spayed

**Figure 3.**
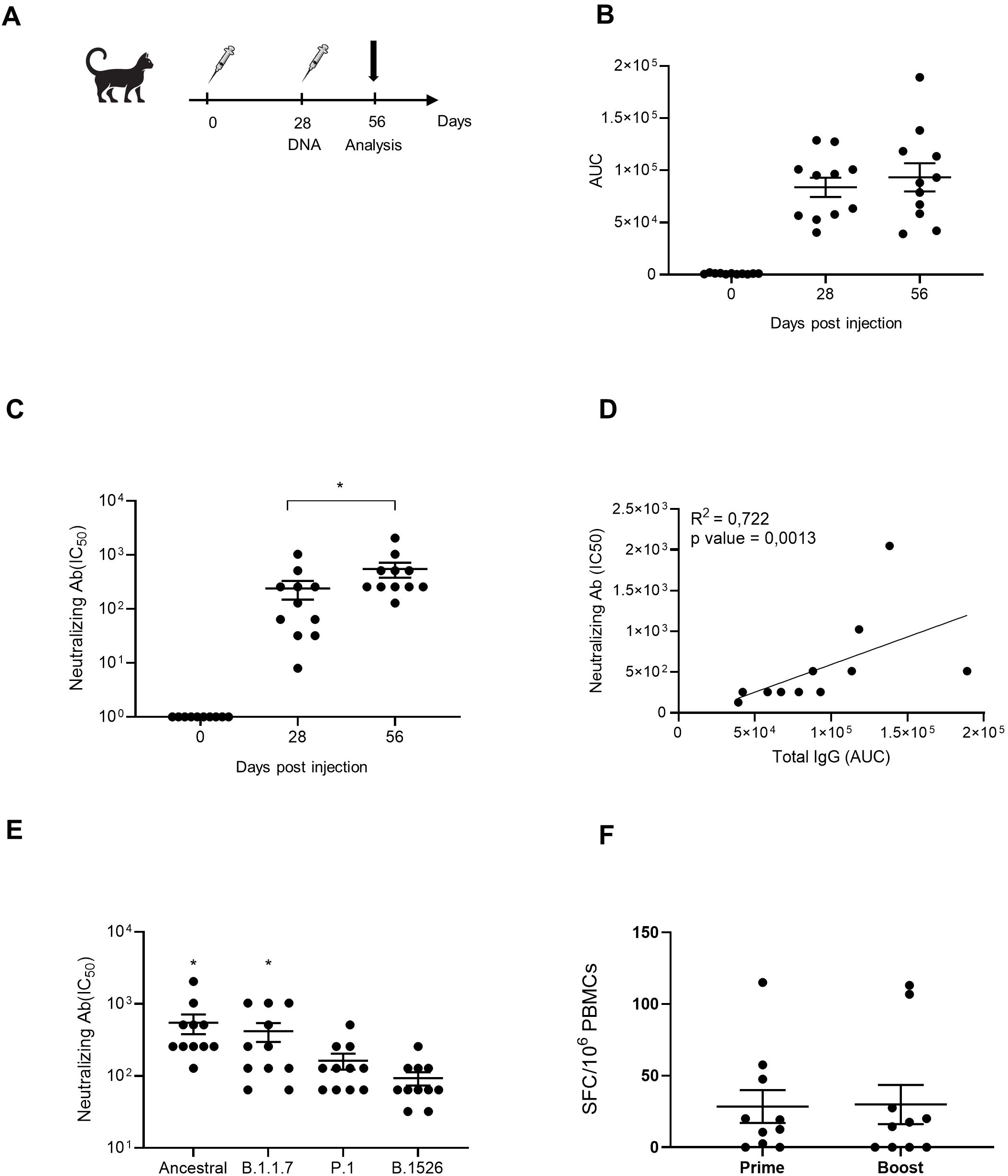
LinDNA immunogenicity in cats. **(A)** Schematic representation of feline study. Eleven client-owned cats, from 1 to 15 years old, were vaccinated by means of i.m. electroporation, with 1 mg of linDNA (o.5 mg per quadriceps) at day 0 (prime) and day 28 (boost). **(B)** Area under curve (AUC) analysis of serum anti-RBD IgG measured at day 0 (before vaccination), day 28 (after prime) and day 56 (after boost). **(C)** Neutralizing antibody titers in sera collected from vaccinated cats at day 0, 28 and 56, measured through a neutralization assay with infectious SARS-CoV-2 and Vero cells. **(D)** Correlation between IgG AUC values and neutralizing antibody IC_50_ values. **(E)** Neutralizing antibody titers in sera collected at day 56 and measured through a neutralization assay with ancestral SARS-CoV-2 and three variants (B.1.1.7, P.1 and B.1526). **(F)** IFNγ-producing T cells measured by ELISpot assay performed on PBMCs collected from vaccinated cats at day 28 (after prime) and day 56 (after boost). Significance was determined using Mann-Whitney test, *p<0,05 **p< 0.01.

To assess potential side effects connected with DNA-EGT, physical examinations, measurements of body weight and temperature were quantified throughout the entire course of the study. No significant change during the entire course of the study or adverse events were identified. Vaccinated animals were monitored for abnormal values with complete blood count, serum chemistry and urinalysis and no significant abnormalities connected with the immunizations were identified demonstrating that the linDNA vaccine is safe when administered to larger animals.

### linDNA vaccine immunogenicity in feline study

In order to assess linDNA vaccine immunogenicity against RBD domain of SARS-CoV-2 Spike protein in domestic cats, we measured anti-RBD IgG levels both after prime (day 28) and after boost (day 56) by ELISA (**Fig.3B**). A consistent antibody titer was present in each vaccinated subject after prime and then increased or remained at same level after boost. We next sought to analyze whether the RBD-specific antibodies induced by the RBD linear vaccine were able to neutralize ancestral SARS-CoV-2 virus and three variants of concern (B.1.1.7, P.1 and B.1.526). More specifically, neutralizing Ab titers against ancestral virus were detectable after prime and significantly increased after boost immunization (**Fig.3C**). Furthermore, neutralizing antibody titers measured after boost significantly correlated with total IgG titers (**Fig.3D**). Neutralizing titers measured against the three SARS-CoV-2 variants were measured after boost and showed to be consistent as well, although neutralizing titers measured against ancestral virus and B.1.1.7 variants were significantly higher than those detected against P.1 and B.1.526 variants (**Fig.3E**).

Finally, we sought to determine the cellular immune response elicited by linDNA vaccine in 10 out of 11 felines vaccinated with prime-boost regimen, by performing ELISpot assay on PBMCs collected at day 28 (after prime) and day 56 (after boost). As shown in **Fig.3F**, after prime immunization, RBD-specific T cells were detected in 8 out of 10 samples, even if no significant increase was measured in T cell response at day 56 (after boost). Altogether, these results confirm that a PCR-based linDNA vaccine encoding RBD domain of SARS-CoV-2 is safe and able to elicit both humoral and cellular immune response and, most importantly, high titers of neutralizing antibodies one of the key correlates of protection of S-based nucleic acid SARS-CoV-2 vaccines.

## DISCUSSION

SARS-CoV-2 global pandemic, which likely originated from bats, has highlighted the need for a One Health approach to prevent and manage emerging zoonotic diseases. Since its outbreak in late 2019, huge efforts have been made for the development of safe and efficient in eliciting a protective immune response in humans. Nonetheless, as SARS-CoV-2 emerged as a spillover from animals to humans, also spillback to other animal species, especially mammals, has been observed more and more frequently^16,17,18^. After more than two-years of the pandemic, it is quite evident the establishment of multiple domestic animal and wildlife reservoirs of SARS-CoV-2, from felines to minks and deers^4,19,20^. These continuous evolutionary changes and viral adaptation to new species have consequently accelerated the emergence of novel variants, that may eventually affect vaccine efficacy in humans^21^. Reports of human-to-feline transmission of SARS-CoV-2 and of limited airborne transmission among cats have been documented since virus spread^5^, but only very recently cat-to-human transmission of SARS-CoV-2 has been reported^22^. These reports demonstrate the strong zoonotic potential of SARS-CoV-2 and underline the urgent need to halt reverse zoonosis from animals to humans by adopting a One Health approach.

To this aim, here we report about a Phase I/II clinical study of a nucleic acid-based vaccine, consisting of a PCR-based linDNA, delivered by electroporation in domestic cats. Besides being the only genetic vaccination platform already approved for veterinary applications, DNA-based vaccines show numerous advantages, such as fast and scalable manufacturing, easy adaptation to new genetic viral sequences, and differently from other genetic vaccines, long-term stability at room temperature. Moreover, to increase DNA uptake, we adopted EGT technology for intramuscular delivery, a methodology extensively studied previously^23^. Besides this, our vaccine is based on PCR-based linDNA amplicons, an innovative cell-free vaccination platform that has already proven its efficacy in eliciting in vivo immune responses^13^.

Preclinical results have shown that a linDNA vaccine encoding the RBD domain of SARS-CoV-2 is immunogenic in mice when administered in a prime-boost vaccination schedule, by eliciting consistent humoral and cellular immune responses. When administered to cats following a prime-boost vaccination schedule, linDNA vaccine proved to be safe, without any adverse event. Importantly, it showed to be immunogenic by eliciting both binding and neutralizing antibodies, the latter essential to protect from live virus infection, soon after prime, against not only the ancestral SARS-CoV-2 Spike RBD but also against three SARS-CoV-2 variants (B.1.1.7, P.1 and B.1526). Also, a consistent T cell response specific for RBD was detected soon after prime. Further studies are warranted to demonstrate linDNA vaccine efficacy in protecting larger animals from virus challenge, even though feline data encourage further investigation. In summary, linDNA vaccine against SARS-CoV-2 is a highly efficient vaccination platform capable of inducing robust, protective neutralizing antibody and T cell responses in felines and potentially other susceptible animal species.

Importantly, vaccination of susceptible animals could protect them against SARS-CoV-2, but may also interrupt the chain of animal-to-animal and even animal-to-human transmission, thus lowering the risk of emergence of novel SARS-CoV-2 variants with increased virulence in humans. Therefore, One Health strategy should be considered to control the circulation of SARS-CoV-2 in all possible susceptible animals and humans via immunization.

## METHODS

### Synthetic genes and linDNA amplicon constructs

The synthesis and codon optimization analysis of a cDNA encoding the RBD region of SARS-CoV-2 protein S has been performed at Genscript (China). DNA plasmid encoding RBD has been described elsewhere^11^. Briefly, synthetic codon-optimized RBD design took into account codon usage bias, GC content, CpG dinucleotides content, mRNA secondary structure, cryptic splicing sites, premature PolyA sites, internal chi sites and ribosomal binding sites, negative CpG islands, RNA instability motif (ARE), repeat sequences (direct repeat, reverse repeat, and Dyad repeat) and restriction sites that may interfere with cloning. In addition, to improve translational initiation and performance, Kozak and Shine-Dalgarno Sequences were inserted into the synthetic gene. To increase the efficiency of translational termination, two consecutive stop codons were inserted at the end of cDNA. For the construction of RBD encoding DNA plasmid, the cDNA was amplified via PCR by using sequence-specific primers and directionally cloned into the linearized pTK1A-TPA vector by enzymatic restriction PacI/NotI. DNA amplicon construct encoding RBD was synthetized as previously described^13^. Briefly, for the phosphothioate-modified amplicons, a sulfur atom substitutes the non-bridging oxygen in the phosphate backbone of the oligonucleotide. In addition, the five (5’) terminal bases of both the forward and reverse primers were modified to increase DNA amplicon stability^14^. DNA amplification via PCR was performed using a large-scale PCR system using Q5 polymerase (New England BioLabs, USA) and resulted in 2780 bp linDNA amplicon expression cassettes. The PCR mixture consisted of 1X buffer with 0.8mM dNTP, 0.5μM of forward and reverse primers, polymerase 1u/100ul, and plasmid template at 40ng/mL. The PCR reactions were subjected to initial denaturation of 98°C for 90 seconds, and subsequent 30 cycles of 97°C for 80 seconds and 70°C for 4.5 minutes; and followed by final extension at 72°C for 3 minutes. As for purification, the target dsDNA from the PCR amplification was first concentrated by ethanol precipitation and then purified on an Akta Pure 150 FPLC instrument (GE Healthcare, USA) with a GE HiPrep 26/60 Sephacryl S-500 High Resolution size exclusion column and 0.3M NaCl running buffer. The purified DNA solution was ethanol precipitated again and resuspended with phosphate buffered saline (PBS) solution to a final concentration of 1mg/mL± 10% and sterile filtered with a 0.22μm PES membrane. Further analytical characterization of the linDNA amplicons was performed using NanoDrop (Thermo Fisher, USA), 2100 Bioanalyzer, and Alliance HPLC System (Waters, USA). Sanger sequencing of the linDNA expression cassette showed no sequence error as compared to the plasmid DNA template sequence. The linDNA expression cassettes were then lyophilized in a VirTis Genesis Pilot system. All removable internal components of the system were autoclaved. All interior surfaces were wipe sterilized with actril and swabbed for confirmation of sterility by micro testing (plating). The purified sterile linDNA solution was aseptically filled into the sample vials (2 mL glass amber 15×32 mm with 13mm crimp) at predetermined volume, and a 13 mm 2-leg stoppers were applied to the mouth of each vial. Stoppered vials were then placed into an autoclaved bag and placed in a −20 ± 5 °C freezer for 18 hours preceding lyophilization. After the lyophilization program was run, the vials were stoppered under vacuum, and West 13 mm smooth vial caps were applied and crimped manually.

### Transient expression of RBD protein and Western Blotting

HEK293 cells were transiently transfected with SARS-CoV-2 RBD expression vectors, both plasmid and PCR-based linDNA amplicon, using Lipofectamine 2000 Transfection reagent (Thermo Fisher Scientific, USA). Two days later, cells were pelleted and lysed in RIPA buffer (Thermo Fisher Scientific). Cell lysates and supernatants were separated by SDS-PAGE and transferred to nitrocellulose membranes. Immunoblotting was performed by using SARS-CoV2 Spike S1 Subunit primary Antibody (Sino Biological) diluted 1:1000 in 5% milk - 0,05% PBS-Tween20. Chemiluminescence detection was performed by using the ECL™ Prime Western Blotting System (Cytiva, Merck) and acquired by ChemiDoc Imaging System (Bio-Rad).

### Mice

For immunogenicity assessment of linDNA Covid-eVax, 6-8 weeks old BALB/c (H-2^d^) mice (Envigo, Italy) were administered with two doses (prime-boost) vaccination schedule, at days 0 and 28, with 20μg of both plasmid and linear Covid-eVax, delivered i.m. by electroporation as previously described^24,25^. At different time points, antibody and cell mediated immune response were analyzed. Mice were housed according to national legislation and kept in standard conditions according to Evvivax ethical committee approval. All the *in vivo* experimental procedures were approved by the local animal ethics council.

### Feline study design and immunization

From February to June 2021, eleven client-owned cats, aged 1-15, both males and females, SARS-CoV-2 tested negative, otherwise healthy with no clinical disease or mitigating, underlying pathologies, were enrolled in a randomized Phase I/II trial. For each cat, the veterinary staff of the Veterinary Oncology Services (VOS, NY, USA) performed a full initial clinical examination and two doses (prime-boost at days 0 and 28) of 1mg linDNA vaccine were delivered intramuscularly (i.m.) by electroporation. The electroporation was carried out with Vet-ePorator™, a device manufactured by IGEA (Italy) for Evvivax and based on Cliniporator® Technology. Electrical conditions consisted of 8 square unipolar pulses at 110 V, at an interval of 120 ms. The pulse length was 20 ms/phase with a frequency of 8 Hz. Each animal was examined pre-operatively prior to each treatment and anesthetized to carry out the electroporation with a short anesthetic protocol. The entire electroporation procedure, including anesthesia, lasted less than 10 minutes. A 22g IV catheter was placed in the cephalic vein prior to induction with dexmedetomidine hydrochloride followed by propofol to effect allowing placement of 3.5 endotracheal tube. Oxygen was administered together with isoflurane, when needed to provide an even plane of anesthesia. Monitoring for blood pressure, heart rate, pulse oximetry, electrocardiogram and temperature was consistent during the procedure. The patient was positioned in dorsal recumbency and in both regions of the tibialis cranialis muscle, a 1.5cm squared area was clipped and prepared. 0.5 ml of linDNA were administered respectively into each muscle and followed with placement of the N-10 4B electrode to provide the electroporation needed for electro-gene-transfer as confirmed by a 2 second display of muscle reflex in the representative leg.

Blood was collected at indicated time points and serum was frozen for further analysis. An informed consent consistent with the requirements of USDA APHIS relating to the study of an unlicensed veterinary biological product was distributed and approved by all cat owners. Each researcher tested negative via PCR during each administration.

### Laboratory tests

Complete blood count, serum biochemical profile of 12 analytes (including total protein, albumin, urea, creatinine, alanine aminotransferase, aspartate 1aminotransferase, alkaline phosphatase, gamma-glutamyltransferase, calcium, phosphorus, iron, cholesterol, glucose, sodium, potassium in serum) and urinalysis were obtained from each patient.

### ELISpot assays

The T cell ELISpot for mouse IFNγ was performed as previously described^26^. A pool of RBD peptides (consisting of 132 out of the 338 peptides covering the whole Spike protein, JPT) was used for PBMC stimulation. Mouse ELISpot assay was performed by stimulating splenocytes, collected at cull from plasmid and linear vaccinated mice, for 20h with RBD peptide pool. Feline ELISpot assay was performed by stimulating PBMCs, collected at days 0 (before prime), 28 (before boost) and 56 (after boost), for 20h with RBD peptide pool, following manufacturer’s instructions (Mabtech).

### Antibody detection assays

Both in murine and in feline study, antibody titration was performed on sera, collected at different time-points. The ELISA plates were functionalized by coating with the RBD-6xHis protein at a concentration of 1 μg/ml and incubated about 18 hours at 4°C. Subsequently the plates were blocked with 3% BSA / 0.05% Tween-20 / PBS for 1 hour at room temperature and then the excess solution was eliminated. Sera were then added at a dilution of 1/300 and diluted 1:3 up to 1/218,700, in duplicate, and the plates incubated overnight at 4°C. After a double wash with 0.05% Tween-20 / PBS, the secondary anti-murine IgG or anti-feline IgG conjugated with alkaline phosphatase was added and the plates were incubated for 1 hour at room temperature. After a double wash with 0.05% Tween-20/PBS, the binding of the secondary was detected by adding the substrate for alkaline phosphatase and measuring the absorbance at 405nm by means of an ELISA reader after incubation for 2 hours.

Virus neutralization assays were performed at the Animal Health Diagnostic Center at Cornell University (NY, USA). Heat inactivated serum samples were serially diluted (1:8-1:2048) and incubated with 100-200 TCID50 of SARS-CoV-2 (ancestral/D614G, B.1.1.7, P.1 or B.1.526) in 96-well plates. After 1 h incubation at 37^°^C, a suspension of Vero-E6 cells was added to each well of the test plates and incubated at 37^°^C with 5% CO2 for 72 h. Viral cytopathic effect were monitored after incubation and neutralization titers determined as the reciprocal of the highest dilution of serum capable of completely blocking viral replication^27,28^.

### Statistical analyses

Statistical analyses were performed with GraphPad Prism software version 8 (GraphPad). *n* represents individual mice analyzed per experiments. Error bars indicate the standard error of the mean (SEM). We used Mann-Whitney U-tests to compare two groups with non-normally distributed continuous variables. Significance is indicated as follows: *p<0.05; **p<0.01. Comparisons are not statistically significant unless indicated.

## ACKNOWLEDGMENTS

Veterinary Oncology Services would like to acknowledge Michelle Barreto for her help with feline patients and procedure during the study.

## AUTHOR CONTRIBUTIONS

JAI and ES enrolled cats and performed the clinical trial; AC, ES, LL, MC, EP performed experiments in mice and immunological assays on feline samples; DGD performed neutralization assays; AC, FP and LA analyzed and interpreted data; AC prepared the figures; LA provided funding, conceptual advice and edited the manuscript; AC coordinated the study and wrote the paper; BV produced the amplicons; JH and CS provided conceptual advice, coordinated the study and edited the manuscript.

## ETHICS APPROVAL

All animal experiments were performed according to the guidelines for the care and use of laboratory animals and were approved by the ethical committee of the Italian Ministry of Health, with authorization #1166/2020-PR. An informed consent consistent with the requirements of USDA APHIS relating to the study of an unlicensed veterinary biological product was distributed and approved by each cat owner.

## DATA AND MATERIALS AVAILABILITY

All data and materials are available in the main text.

